# Detection of glioma and prognostic subtypes by non-invasive circulating cell-free DNA methylation markers

**DOI:** 10.1101/601245

**Authors:** H Noushmehr, TS Sabedot, TM Malta, K Nelson, J Snyder, M Wells, A deCarvalho, A Mukherjee, D Chitale, M Mosella, K Asmaro, A Robin, M Rosenblum, T Mikkelsen, J Rock, LM Poisson, I Lee, T Walbert, S Kalkanis, AV Castro

## Abstract

Genome-wide DNA methylation profiling has shown that epigenetic abnormalities are biologically important in glioma and can be used to classify these tumors into distinct prognostic groups. Thus far, DNA profiling has required surgically resected glioma tissue; however, gliomas release tumoral material into biofluids, such as blood and cerebrospinal fluid, providing an opportunity for a minimally invasive testing. While prior studies have shown that genetic and epigenetic markers can be detected in blood or cerebrospinal fluid (e.g., liquid biopsy [LB]), there has been low sensitivity for tumor-specific markers. We hypothesize that the low sensitivity is due to the targeted assay methods. Therefore, we profiled the genome-wide CpG methylation levels in DNA of tumor tissue and cell-free DNA in serum of glioma patients, to identify non-invasive epigenetic LB (eLB) markers in the serum that reflect the characteristics of the tumor tissue. From the epigenetic profiles of serum from patients diagnosed with glioma (N=15 *IDH* mutant and N=7 *IDH* wildtype) and with epilepsy (N=3), we defined glioma-specific and *IDH*-specific eLB signatures (Glioma-eLB and *IDH*-eLB, respectively). The epigenetic profiles of the matched tissue demonstrate that these eLB signatures reflected the signature of the tumor. Through cross-validation we show that Glioma-eLB can accurately predict a patient’s glioma from those with other neoplasias (N=6 Colon; N=14 Pituitary; N=3 Breast; N=4 Lung), non-neoplastic immunological conditions (N=22 sepsis; N=9 pancreatic islet transplantation), and from healthy individuals (sensitivity: 98%; specificity: 99%). Finally, *IDH*-eLB includes promoter methylated markers associated with genes known to be involved in glioma tumorigenesis (*PVT1* and *CXCR6*). The application of the non-invasive eLB signature discovered in this study has the potential to complement the standard of care for patients harboring glioma.

## INTRODUCTION

Gliomas are a heterogenous group of intracranial tumors that are constantly evolving, generally recur, and frequently progress to more malignant subtypes. Recently, genomic and epigenomic alterations have defined subtypes of glioma (e.g., *IDH* mutation, 1p19q chromosomal deletion, and Glioma-CpG Island Methylator Phenotype [G-CIMP]) with distinct prognostic outcomes ^1–6^. Currently, this molecular diagnosis and classification, which guides clinical management, is dependent on tissue profiling obtained by invasive surgical approaches (tissue biopsy or excision). However, this surgery-centered approach does not allow serial tissue evaluation to capture the dynamic molecular evolution of these tumors, may not be feasible in surgically inaccessible tumors, requires the risk of an invasive procedure in an often comorbid population, and may delay the diagnosis of this disease to later stages due to procedure risks and diagnostic sensitivity. MRI is a relevant non-invasive approach to diagnose and follow patients with glioma; however, limitations remain for differential diagnosis (e.g., lymphoma), detection of minimal or remnant tumoral burden, and in distinguishing progression from pseudo-progression caused by radiation-induced necrosis or treatments such as immunotherapy ^7,8^. In addition, serial assessments may be costly and cumbersome procedures for patients. Therefore, the discovery of a minimally or non-invasive approach that allow earlier identification of sensitive and specific molecular biomarkers that reflect tumor burden and its dynamic evolution in real-time is desirable. An approach that meets the above criteria is liquid biopsy (LB) of biofluids (e.g., blood, and cerebrospinal fluid [CSF]) which detect materials shed by the tumors such as circulating tumor cells and genomic specimens (e.g., circulating tumor DNA) ^9,10^.

In the past decade, investigation of the diagnostic, prognostic and predictive applications of LB throughout a patient’s disease course has been feasible in many tumors ^11–16^. For instance, in central nervous system (CNS) neoplasms, including gliomas, CSF has been a relevant source of molecular markers ^10,17–29^ and can be used to track the glioma tumor evolution ^30^. However, obtaining CSF is an invasive, complex and risky procedure which can cause, for instance, brainstem herniation due to increased intracranial pressure and/or risk of bleeding due to thrombocytopenia caused by chemotherapy in patients harboring CNS tumors. In addition, serial assessment of CSF markers throughout a person’s disease follow-up as standard clinical management raises additional concerns of patient compliance, impact on quality of life, feasibility to perform test (e.g., patients on anticoagulation or those unable to lie flat) and additional risks secondary to repeated use of this procedure. In contrast, blood LB is minimally invasive, quick and feasible to perform longitudinally; however, one limitation of blood-derived LB is the dismal and often low yield of molecular material released into the blood by CNS tumors (likely due to the blood brain barrier) which may hinder the detection of molecular features, such as specific and rare tumor mutations or novel and clinically relevant molecular markers shed by the tumor ^10,21,31,32^ To overcome these limitations, the performance of a more comprehensive “omics” approach (e.g., DNA methylomic or genomic) has been proposed ^10,33^ and implemented for certain cancer types^31,34^ Although genomic LB associated with gliomas has been performed ^10,26^, the anticipated success of highly specific LB markers has been hampered by the low mutation frequency of the associated glioma tumor (0.5-2.6%) ^3,35^. In addition, genetic alterations are generally site specific (e.g., frame-shift, point mutation) which may limit their detection in the fragmented DNA released into the circulation ^36^. The low likelihood of detecting at least one of the 75 significantly mutated genes (somatic)^3^ associated with glioma in any cell-free DNA assay renders the feasibility of genomic-specific LB a challenge. On the other hand, DNA methylation is a stable marker that is tissue specific, clinically relevant to gliomas, and altered in large regions of the genome. Thus, DNA methylome profiling is an attractive approach for the identification of diagnostic, prognostic and predictive markers in LB ^34,37^ In the tumoral tissue, genome-wide DNA methylation profiling has shown that epigenetic abnormalities play important biological and clinical roles in CNS tumors, particularly in gliomas ^3,38–41^. For instance, G-CIMP is a subset of glioma with extensive epigenomic alterations that confers a stronger prognostic value than age, grade and histology ^1,3,41^. However, the relevance of comprehensive DNA methylation profiling in the blood-based circulating free tumor DNA of patients with gliomas has not been explored. Herein, we hypothesized that genome-wide methylome profiling is a useful approach to identify methylome-specific markers associated with glioma in cfDNA. To address this hypothesis, we profiled the genome-wide CpG methylation landscape of matching serum and tissue from 22 patients diagnosed with gliomas (N=15 *IDH* mutant and N=7 *IDH* wildtype) and from 3 patients with a non-neoplastic brain disease (i.e., epilepsy). We identified a set of epigenetic signatures in the serum LB (herein referred to as eLB) that resembles the tissue epigenomic landscape associated with glioma. We showed that the eLB could differentiate glioma from non-tumoral brain tissue and stratify gliomas based on prognostic class (e.g., *IDH* mutation status). We further observed that the specificity of the eLB allowed accurate discrimination of patients with glioma from patients with tumors of other origins and from patients with immune-related disease states (pancreatic islet transplantation and sepsis). The *IDH*-eLB signature includes promoter methylated markers associated with genes known to be involved in glioma tumorigenesis (e.g., *PVT1* and *CXCR6*). Finally, we propose a novel clinical approach to apply the eLB panels to complement the standard of care in the diagnosis and follow-up. The ability to monitor patients by eLB has the potential to improve the pre- and post-surgical quality of care for patients harboring gliomas.

## RESULTS

### Glioma cell-free DNA methylome

In this study, we selected 22 matching pairs of primary glioma tissue and serum, stored at the Hermelin Brain Tumor Center (HBTC) bank from patients who underwent neurosurgery at the Henry Ford Health System, Detroit, Michigan. Serum was collected immediately prior to dura incision at the time tumor tissue was surgically resected. According to the current World Health Organization (WHO) 2016 criteria for glioma classification, our HBTC cohort comprised 3 grade IV, 11 grade III, and 8 grade II gliomas among which 15 *IDH* mutants (*IDH*mut) and 7 *IDH* wildtypes (*IDH*wt) were included (Table 1, Extended Data Table S1). As expected, we observed a significant overall survival difference between patients with *IDH*mut and *IDH*wt tumors (mean (95% CI): 62.62 months (53.72-73.32) *versus* 26.86 months (8.85-33.6), Extended Data Fig. S1A). Total extracted serum cfDNA quantity, normalized to the genomic size (Genomic Equivalents [GE]/ml, see Methods), showed that patients with glioma had significantly higher serum cfDNA in relation to patients who underwent surgery for epilepsy in the absence of tumor (mean±s.e.: 12268.8±9269.1 *versus* 3777.4±2324.7 GE/ml, respectively, student t-test p-value=0.003216; Fig. 1A). This is consistent with the hypothesis that tumors tend to release more cfDNA. Aligned with the hypothesis that increased cfDNA is associated with aggressive tumors, likely due to brain-blood barrier breakdown ^42,43^, we also observed a trend, albeit not statistically significant, wherein *IDH*wt serum had more cfDNA than *IDH*mut (mean±s.e.:15799.3±8645.4 vs 10621.2±9364.9 GE/ml; respectively, student t-test p-value=0.2255, Extended Data Fig. S1B) suggesting that cfDNA is altered during gliomagenesis and may be released more abundantly in more aggressive subtypes.

**Fig. 1:**
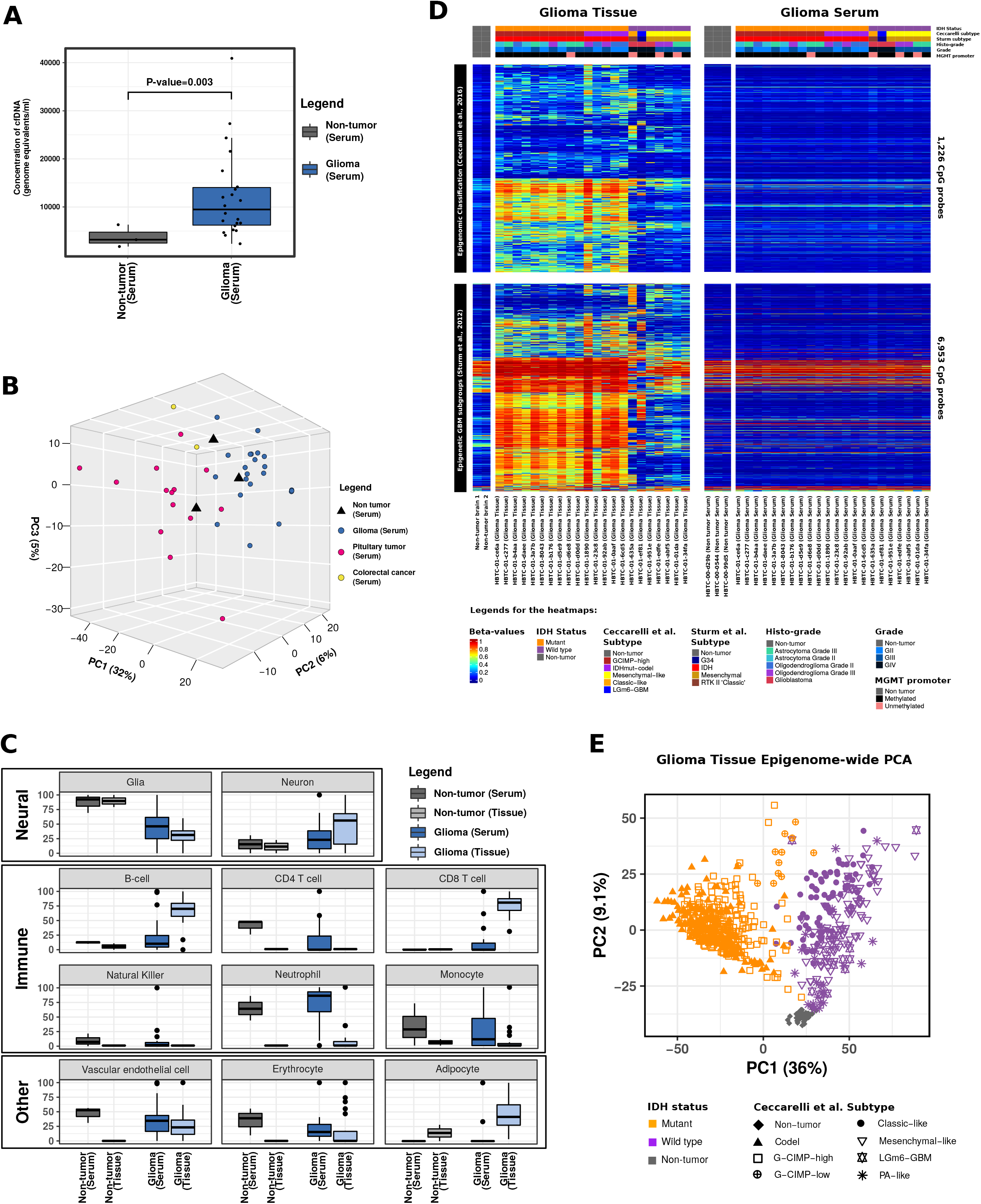
Genome-wide DNA methylation profile of Glioma serum cfDNA. A) Total serum cfDNA concentration normalized to the genomic size (Genomic Equivalents/ml) is represented in two boxplots (non-tumor vs glioma). B) Epigenome-wide cell-free DNA methylation (serum-based) derived from Glioma, Pituitary tumor, Colorectal carcinoma (CRC), and non-tumor patients presented by a similarity method: Principal Components Analysis (PCA). Each dot represents a sample cfDNA methylation (epigenome-wide) and colored based on tissue/tumor of origin. Total percent variance is indicated along all three axes. C) Cell-type CpG methylation-based deconvolution of our patients’ cfDNA methylome is divided into three relevant categories: Neural, Immune and Other cell types. Y-axis represents normalized percent of each cell-type and separately by serum and tissue present in the cfDNA methylome. Eleven sets of boxplots are each divided into two categories; non-tumor (grey) vs glioma (blue). The plots are further divided by tissue (non-tumor-light grey or glioma-light blue) and cfDNA serum (non-tumor-dark grey or glioma-dark blue). D) Published tissue-derived epigenetic signatures from Ceccarelli et al. 2016 (top heatmaps) and Sturm et al. 2012 (bottom heatmaps) are undetectable in Glioma cfDNA methylome. Levels of CpG methylation in our HBTC cohort divided by glioma tissue (left two heatmaps, respectively) vs matching serum cfDNA (right two heatmaps). Columns indicate patients with clinical and molecular annotation tracks listed on top of heatmaps and rows indicates CpG probe. Beta-value (DNA methylation levels) are indicated in the legend from 0 (low CpG methylation) to 1 (high CpG methylation). E) Glioma-tissue epigenome-wide PCA highlights the difference between glioma and non-tumor and between *IDH*mut and *IDH*wt patients. TCGA Pan-Glioma DNA methylation data from Ceccarelli et al. 2016 is represented in a Principal Component Analysis to evaluate similarities genome-wide. The first two principal components are plotted using all available DNA methylation data points (~400,000 CpGs, unfiltered). The variance total percentage is labeled along both axes. Fifty-four total non-glioma tissue (grey) were also included in this analysis to highlight the epigenome-wide difference between glioma and non-tumor. The glioma cohort is further divided by available known IDH status and by our recent epigenomic subtypes.

We performed an epigenome-wide profile of the glioma cfDNA using Infinium Human Methylation 850K (HM850K). Filtering and pre-processing steps were taken to align these data with that of serum methylome data from colorectal cancer (N=2) ^44^ and pituitary adenoma (N=14, unpublished data), as well as from glioma tissue ^3^ (see Methods). Principal component analysis (PCA) showed a distinct separation between gliomas and non-tumor specimens as well as other neoplasms (Fig. 1B). We estimated the source or content of the released cfDNA, and DNA of the matched tissue specimens, by deconvoluting the methylome using primary non-cancer cell-type methylation-based signatures ^45^. We observed that in relation to non-tumor specimens, the methylomes from both serum and tissue of glioma patients tended to present less glial (52% lower proportion on average in tumor compared to non-tumor) and more neuronal cell-related (79% greater proportion on average in tumor compared to non-tumor) epigenetic signatures (Fig. 1C, Extended Data Table S2). Interestingly, for specific signatures related to the immune cells, the serum from glioma patients showed a distinct makeup in relation to their matching tissue and non-tumor serum. For instance, B-cell and CD8 T-cell-related signatures in the glioma-patient serum were lower than in the tissue, but glial-related specimens were higher than the associated non-tumor specimens (1.9 and 22 fold increase in means for serum, respectively; 12 and 130 fold increase in means for tissue, respectively; Fig. 1C). On the other hand, CD4 T-cell-related signatures were depleted in the tumor serum in relation to non-tumor serum (0.38 fold decrease) and undetectable in the tumor and non-tumor tissue. Notably, the serum from both non-tumor and glioma patients included signatures associated with higher proportions of neutrophil- and monocyte-related cell types. Together, these results indicate that the cfDNA methylome contains signatures that are specifically related to glioma as well as to the immune system, which may be related to a response to changes in the tumor microenvironment or release of immune invasion from the glioma tumor.

Considering the full methylome profile, we investigated whether glioma subtype-specific epigenetic signatures defined at the tumor tissue level could be detected in the serum of glioma patients. Interestingly, despite evidence that brain tumors release circulating tumor DNA into the blood and that the release mechanism is dependent on a phenotype modification (e.g., mesenchymal glioma subtype) ^46^, the published epigenetic signatures ^3,6^ were undetectable in the serum methylome (Fig. 1D & Extended Data Fig. S1C-F). This is consistent with previous findings using whole-genome sequencing that have shown a low detection of tumor genetic hotspot mutations associated with gliomas in the cfDNA ^10,26^. However, given that we observed epigenome-wide differences in the tissue of glioma patients in relation to other tumor types and non-tumor tissue samples (Fig. 1E), it is plausible that other relevant CpG methylation sites, not previously included in glioma marker panels, could be released into the serum by the brain tumor and, consequently, be detectable *via* analysis of the serum methylome.

### Glioma eLB derived from cell-free DNA in serum can identify patients with glioma

We performed an epigenome-wide differential analysis to identify specific, serum-based, epigenetic markers associated with glioma compared to non-glioma serum methylome, which we named Glioma-eLB (N=1075 differential CpG sites overlapping autosomal chromosomes, Wilcoxon-rank sum test p-value < 0.001, Extended Data Fig. S2A-B). We confirmed the origin of the Glioma-eLB by evaluating the measured DNA methylation for each CpG in the matching glioma tissue. The glioma tissue-derived methylome was analyzed as part of TCGA project using the Illumina HM450K array, a reduced content compared to the HM850K. We identified 567/1075 differential CpGs measured on the HM450K array, of which 384 (68%) of the detectable Glioma-eLB were present in the matching tissue and distinct from serum methylome of other neoplasias (Fig. 2A-B, Extended Data Fig. S2C, Extended Data Table S3). We explored the specificity of the Glioma-eLB signature using independent primary glioma tissue and other non-glioma tumor methylomes and confirmed that glioma serum methylome clustered with primary glioma (Fig. 2C, Extended Data Fig. S2D), corroborated our initial observation that the eLB measurements reflected the glioma-tissue methylome. To further investigate the content of the Glioma-eLB signature, we analyzed the similarity matrix across available non-cancer cell-type signatures based on methylation (brain-, neural- and immune-associated cell types) and observed that Glioma-eLB was more similar to neutrophils, monocytes, and normal glia- and neuronal-cell-related signatures than any other normal cell types originating from different cell lineages (Fig. 2D, Extended Data Fig. S2C). Interestingly, the Glioma-eLB signature segregated with non-tumor serum that in turn clustered with brain and glial signatures, suggesting that Glioma-eLB signature also captures tissue-of-origin signatures. Overall, we found that non-invasive serum-derived Glioma-eLB signature is detectable and may reflect the expected heterogeneous cell population present in glioma tissue.

**Fig. 2:**
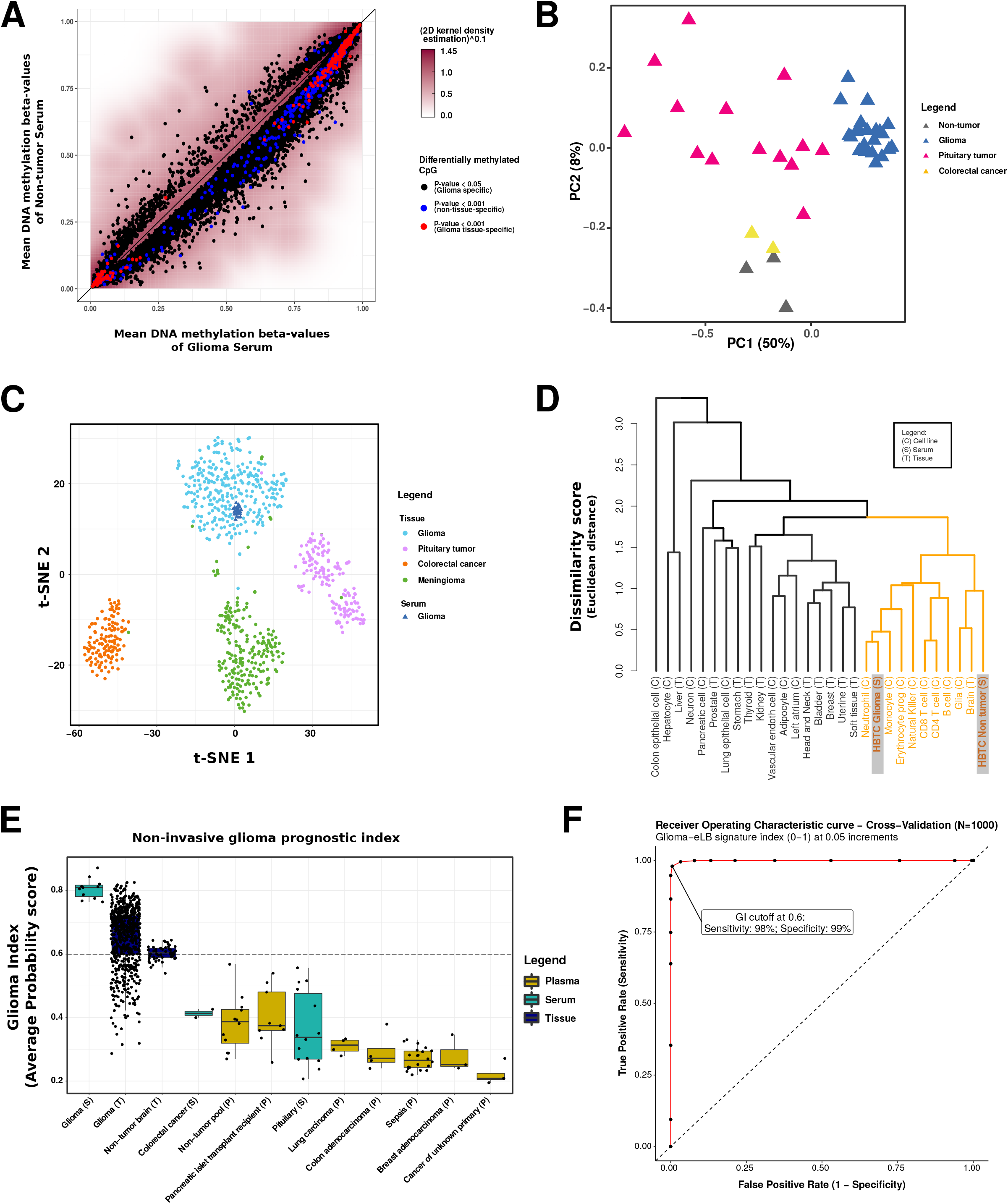
Identification of Glioma-specific Epigenetic Liquid Biopsy (eLB) as a Diagnostic Marker for Gliomas. A) Epigenome-wide mean DNA methylation across our patient cohort’s serum cfDNA methylation (y-axis: three nontumor samples and x-axis: 22 glioma-patient-derived serum samples). Non-significant CpG methylation probes (p-value > 5%) are condensed into a density heatmap by calculating the 2D kernel density estimation to the power of 0.1. Identified cfDNA methylation signatures associated with glioma patients (N = 1,075) are selected by different p-values (black, p-value < 0.05; blue, and red, p-value < 0.001). Glioma-eLB signatures (p-value < 0.001) are further divided into CpGs that are measurable (Glioma tissue specific, red) or not measurable in the matching glioma tissue (Non-tissue specific, blue). B) Principal Component Analysis using the Glioma-eLB signatures as input. Serum methylome from glioma, pituitary tumors, CRC and non-tumor samples are represented. C) Glioma-specific tissue-matching eLB (N=384) was used to subset the published primary tumor tissue DNA methylation and using t-SNE (t-distributed stochastic neighbour embedding) dimensionality reduction to visualize the similarities of each sample. As expected, each primary tumor type (circles) clusters with its known cell-of-origin. Serum cfDNA methylation of our patients cohort (triangles) clusters with the primary glioma tissue DNA methylation profiles. D) Dendrogram of non-tumor cell types in comparison to Glioma-eLB (N=384). Based on the mean DNA methylation across each cell type, this dendrogram shows that glioma serum cfDNA clusters with relevant immune cell-types along with glial-derived cells and bulk brain (non-tumor) samples. E) Machine learning (ML) application (Random Forest) using our defined Glioma tissue-specific eLB to classify tumors and available cfDNA methylation (serum or plasma) derived from tumor patients, patients with metastasis of unknown primary, non-tumor conditions (e.g., sepsis, pancreatic islet transplantation recipient) and non-tumor/non-diseased cell-free DNA. Y-axis represents the ML similarity index based on Glioma-eLB signatures averaged across 1000 iterations. Zero indicates low probability of a glioma, while 1 indicates high probability of a sample being a glioma. Dash line indicates cutoff to determine glioma classification. F) Receiver operating characteristic curve derived from the average across 1000 cross validation. Specificity and sensitivity calculated at 0.05 increments (N=21) from 0 to 1 Glioma-eLB signature index.

Next we annotated the genomic location of the Glioma-eLB. We observed 167 CpG probes (see Extended Data Table S3) that overlapped with known promoters; 64 were hypermethylated and not significantly enriched in promoters (OR=0.92, 95% CI: [0.69, 1.21], chi-square test p-value=0.564) while 103 were hypomethylated and this observation was enriched in promoters (OR=2.36, 95% CI: [1.84, 3.04], chi-square test p-value=4.49E-12) above expected distribution for the methylation platform (Extended Data Table S4). The lack of matching gene expression data to the normal brain tissue DNA methylation limited our ability to investigate the biological context of this Glioma-eLB signature.

Given the specificity of the detectable Glioma-eLB, we developed a machine learning (ML) model to predict the presence of glioma. To determine the robustness of our eLB signature, we applied a cross-validation method as follows: First, we redefined a new set of Glioma-eLB relative to non-tumor controls using 11 randomly selected cases from the initial cohort (N=22). Next, we trained a ML model (RandomForest) using the Glioma-eLB on the training set. This provided us with a Glioma-eLB Index (GI), which estimates the probability that a sample is likely a glioma (high GI or close to 1) or non-glioma (low GI or close to 0). We evaluated the performance of the ML model and GI by applying it to the serum methylome of gliomas of the test set (serum specimens left out of the training set, N=11), as well as to other serum cfDNA and plasma methylomes from patients with non-tumor conditions (sepsis, pancreatic islet transplant recipient) and other neoplasias (colorectal, breast, lung, pituitary and cancers of unknown primary). To assess the stability of the GI development method, we replicated the test-set selection, ML generation, and application steps 1000 times. The averaged GI for each of the 1000 models revealed glioma tissue and serum (test set) methylome samples carried the highest GI (>0.5), whereas the plasma and/or serum of other non-glioma tumors (pituitary tumor, CRC, breast carcinoma, etc.) carried a lower GI (<0.5) (Fig. 2E, Extended Data Table S5). Notably, inflammatory conditions such as sepsis which carry a higher immune response, were not classified as glioma (Fig. 2E), suggesting that despite the close association between Glioma-eLB and immune cell signatures (Fig. 2D), the immune response captured in the cfDNA is specific to gliomas. We evaluated each model’s GI value at increments of 0.05 and determined that a GI of 0.60 accurately (mean±s.e.: 99.3%±0.000278) classified a glioma with 98% sensitivity and 99% specificity (±s.e.: 0.001507 and 0.0002612, respectively, Fig. 2F, Extended Data Table S5). In summary, we conclude that a classification model based on the Glioma-eLB signature can predict a patient’s glioma-like status from other neoplasias, non-cancer diseases or conditions using the methylome profiles identified from a serum sample.

### Identification of prognostic glioma classes by non-invasive eLB

Somatic mutation in one of the isocitrate dehydrogenase genes (*IDH1, IDH2*, [IDH]) is a prognostic marker for adult glioma (WHO Grade II-IV) which is traditionally identified from excised glioma tissue. We sought to define *IDH* mutation status by analyzing the cfDNA methylation data of serum from *IDH*wt and *IDH*mut tumors, using the same approach used to define the Glioma-eLB. Since our cohort carried a low sample size for the *IDH*mut 1p19q codeletion (N=5) and *IDH*mut 1p19q intact (N=10), we combined these two good prognostic subtypes into one class, *IDH*mut (N=15). Applying a supervised method and restricting our analysis to CpGs within autosomal chromosomes (chrom 1-22), we identified 2647 *IDH*-eLB that distinguished *IDH*mut from *IDH*wt gliomas (Fig. 3A, Extended Data Table S6) further refined by selecting those with a similar methylation pattern in the matching tissue (CpGs overlap HM450K = 1525/2647) which generated specific *IDH*mut-eLB and *IDH*wt-eLB signatures (N=114/1525, 7.5% and N=124/1525, 8%, respectively, Fig. 3A, Extended Data Fig. S3A). Harnessing the matching tissue methylome as well as pan-glioma methylome data from adult patients, we observed that the *IDH*-specific eLB discriminates the two *IDH* subtypes at the primary tissue level and the respective *IDH* serum methylome (*IDH*wt and *IDH*mut) clusters with the respective tissue subtype (Fig. 3B-C, Extended Data Fig. S3B-D) corroborating the specificity of the identified *IDH*-eLB.

**Fig. 3:**
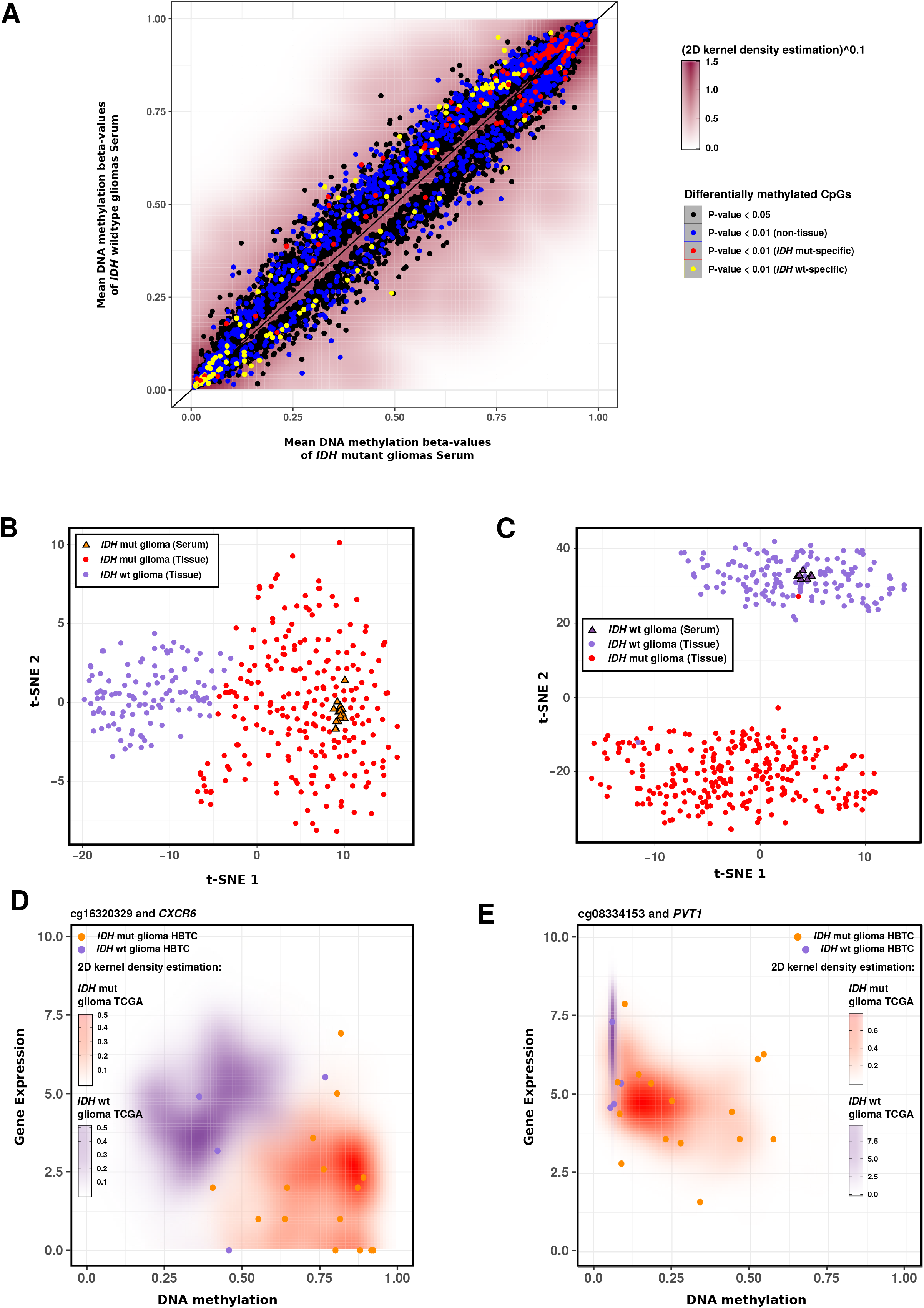
IDH Specific eLB Prognostic Markers. A) Mean DNA methylation of 750,000 CpG across 15 *IDH* mut-patient derived serum (x-axis) vs 7 *IDH* wt-patient derived serum (y-axis). Non-significant CpG methylation probes are condensed into a density heatmap by calculating the 2D kernel density estimation to the power of 0.1. Proposed prognostic glioma-specific eLB (N=1,075) selected by p-values are represented by colored dots (black, p-value < 0.05; blue, red and yellow, p-value < 0.01). Similarities across selected tumor tissue and serum, using *IDH* mut-tissue specific eLB levels (red circles) and *IDH* wt- tissue specific eLB levels (yellow circles). B-C) Similarities across Pan-Glioma tissue (N = 259 *IDH* mut and 160 *IDH* wt) and IDH glioma cfDNA methylation (N = 15 *IDH* mut, 7 *IDH* wt), using *IDH* mut-tissue specific eLB signatures B) and *IDH* wt-tissue specific eLB signatures C) as input in a t-SNE analysis to visualize the similarities by sample. Circles represent primary tissue and triangles represents tumor serum cfDNA. Red indicates *IDH* mut and purple indicates *IDH* wt. D-E) scatter plot between DNA methylation (x-axis) and Gene expression (y-axis) for all pan-glioma primary tumor tissue. 2D kernel density indicates all glioma samples divided by *IDH* status (purple = *IDH* wt, orange = *IDH* mut). Circles indicate the HBTC primary tumor tissue DNA methylation and expression values. D) DNA methylation and expression scatter plot for promoter CpG associated with *CXCR6*. E) DNA methylation and expression scatter plot for promoter CpG associated with *PVT1*.

We then investigated the potential functional or biological role of the *IDH*-eLB by analyzing the methylome and transcriptome from the matching glioma tissue (see Methods). *IDH*wt specific eLB overlapping CpG islands, shores and open seas were significantly enriched or depleted when compared to the expected distribution set by the platform (chi-square test p-value 3.7E-04 enriched, 7.8E-05 enriched, 8.9E-05 depleted, respectively, Extended Data Fig. S3E). We identified 28 *IDH*-eLB-specific signatures linked to a gene promoter (Table 2) of which, 14 transcripts were differentially expressed in *IDH*mut vs *IDH*wt tissues. Ten out of 14 transcripts were inversely expressed in relation with the promoter methylation state (i.e. hypermethylated promoter and down-regulated expression or vice-versa) (summarized in Table 2, Extended Data Fig. S4A). For instance, *CXCR6* is a chemokine related to *CSCL16*, that is overexpressed in glioma and associated with poor prognosis ^47–49^. *CXCR6* expression has been reported as a predictor of recurrence and survival in hepatocellular carcinoma, in addition to intratumoral neutrophils ^50^ and interestingly, *CXCR6* knockout glioma mice survived longer^47^. Congruent with these reports, *IDH*-eLB signature include the promoter hypomethylated state of *CXCR6* and this gene is overexpressed in both TCGA and HBTC *IDH*wt glioma tissues (Fig. 3D). *PVT1* is a long noncoding RNA (lncRNA) that when highly expressed is associated with progression and poor prognosis in a pan-cancer cohort from TCGA. *PVT1* overexpression has also been associated with poor response to chemotherapy in gliomas and squamous cell carcinoma of the head and neck ^51–53^. In line with these findings, hypomethylation of the *PVT1* promoter was detected in our *IDH*-eLB in association with overexpression of the correspondent gene in the worst prognostic subtype (*IDH*wt), in both the TCGA and the current cohort samples (Fig. 3E). Altogether, these findings suggest that with the selected *IDH*-eLB were able to differentiate the various prognostic subtypes of glioma by utilizing the serum methylome. Moreover, the associated CpG methylation in promoter genes which carry a biological and prognostic value in gliomas, support the idea that these non-invasive eLB signatures are feasible for detection and specific to glioma tumors.

## DISCUSSION

In a review by Wan et al ^33^, studies to define non-invasive blood-based markers for cancer has mainly focused on identifying circulating tumor DNA (ctDNA) somatic mutations. And as such, the focus has been on obtaining ctDNA from plasma since blood cell lysis during the preparation of serum samples could release DNA from non-cancerous cells and thus dilute any ctDNA markers and complicate the detection of ctDNA ^33^. However, in glioma in particularly, low sensitivity and reliability of extracting any glioma-specific ctDNA from plasma remains a challenge because of the low frequency in somatic mutation and the targeted approach ^36^. Since epigenetic reflect the cell-of-origin ^54^ and in glioma, somatic DNA methylation aberrations are widespread ^1,3,39,41^, we focused on profiling the DNA methylation of the released DNA to detect glioma status (Glioma-eLB) and associated prognostic subtypes (*IDH*-eLB) by blood (summarized in Fig. 4A). In addition, epigenetic profiling has the advantage of providing information about the tumor microenvironment, which usually lacks somatic mutations ^33^. Deconvolution analysis allowed us to estimate that the Glioma-eLB were associated with signatures from immune, neuronal, and glial cells; however, specific contribution of the individual cells that comprise glioma could not be fully assessed as the information regarding the methylome of the individual tumor cells is currently lacking. The signatures that characterize Glioma-eLB are evidence in favor of leakage of the tumor microenvironment, or from the tumor itself, as well as of a systemic immune response due to the presence of the glioma ^55^. The application of a variety of integrative approaches using matching methylome and transcriptome of the primary tumor as well as available serum/plasma methylome from other non-glioma patients highlighted the sensitivity and specificity of the detected Glioma-eLB. For instance, the ML algorithm using the Glioma-eLB as the input correctly classified glioma tissues as such, in contrast to the serum or tissue of other tumors and non-neoplastic conditions that were not classified as gliomas (Fig. 2E). Despite Glioma-eLB comprising immune-cell-signatures, patients with pro-inflammatory processes, such as sepsis, were not classified as glioma suggesting that Glioma-eLB signatures related to the immune cells were related to the presence of glioma (Fig. 2E). Many previous studies had limited success in detecting well known prognostic markers, currently used in clinical practice and accrued from tissue analysis (such as *MGMT* status, *IDH, PTEN, EGFR*, etc.) ^10,56,57^; however, our holistic approach of unbiased screening allowed us to detect *IDH*-eLB overlapping promoters of genes which transcripts have been well characterized as having a functional role in glioma tumorigenesis (e.g., *PVT1, LOXL, CXCR6*, etc.) even though they are not currently utilized as part of the clinical treatment strategy. These results also highlight that the *IDH*-eLB signatures are capturing glioma-tissue specific markers.

**Fig. 4:**
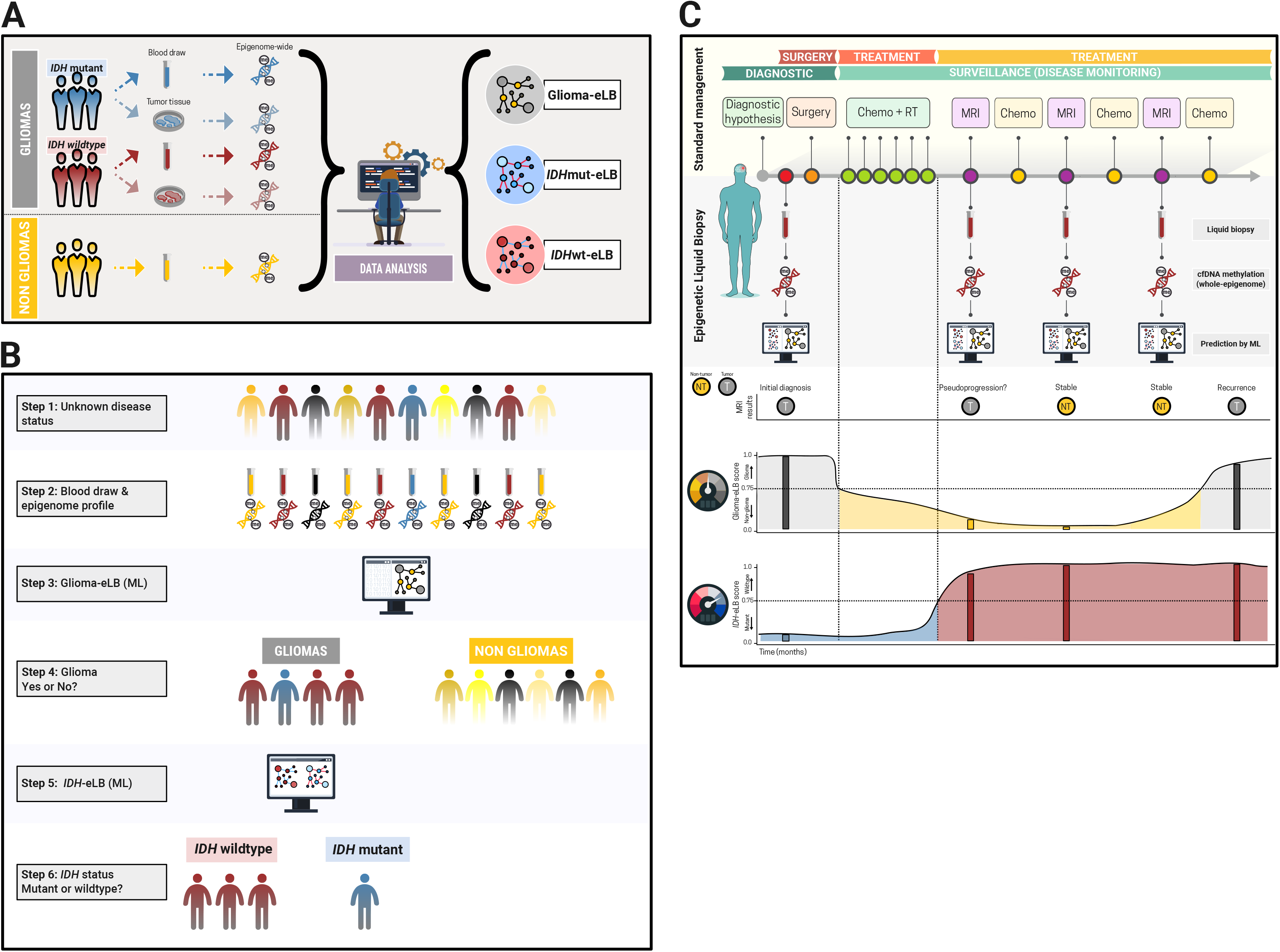
Proposed clinical application of Glioma- and *IDH* eLB. Real time diagnosis and, surveillance of early tumor progression or recurrence. A) Steps used to generate the eLB and the application of a machine learning (ML) model to predict glioma and glioma subtypes. B) At specific intervals, currently determined by magnetic resonance imaging (MRI) visits, patients’ whole-blood is collected and serum/plasma is immediately processed. cfDNA is isolated and profiled using DNA methylation microarray (profile >700,000 CpGs across the entire human genome) and entered into a ML algorithm to generate a index (Glioma-index). C) According to the predetermined index threshold (0.6, sensitivity: 98%; specificity: 99%), the new sample is classified as glioma or non-glioma and according to *IDH* status (mutant or wildtype). The discovered glioma specific eLB (N = 1,075) can be used to complement current clinical diagnostic and monitoring events. Prognostic *IDH*-eLB could be used at time of diagnosis and for monitoring during active treatment through survivorship care. Complementing the MRI findings, eLB could improve detection, reduce false-positive and increase tumor identification (glioma vs necrosis vs non glioma conditions), assess treatment outcome and help tailor treatment options for patients with glioma. eLB could also foster early detection of glioma progression to improve treatment outcomes. Future clinical trials are needed to evaluate the robustness of our eLB signatures for clinical application.

Once validated in the proper settings (e.g., independent cohort), the potential clinical implications of our findings could pave the way for altering current standard diagnostic measures and therapies for glioblastoma, and possibly other brain tumors. Although still in its infancy, the detection of glioma in the blood using a probability score generated by a ML algorithm based on the methylome profile of the patient could challenge the current paradigm of a high-risk tissue-dependent diagnosis prior to definitive treatment. For instance, the application of this approach to patients with tumors not amenable to a meaningful resection due to comorbidities or neuroanatomical surgical limitation such as deep or eloquent regions in the brain could have a major impact in their management. In such scenarios, patients and physicians could proceed to chemoradiotherapy without tissue diagnosis, negating the morbidity associated with biopsy or subtotal resections that offer limited therapeutic benefit while starting definitive treatment sooner. In our proposed model (Fig. 4B-C) and as described by others, the liquid biopsy could also help differentiate glioma from other conditions (e.g., CNS lymphoma; demyelinating disease or metastatic disease) and solve the dilemma of radiographic confounders, such as pseudoprogression from radiation necrosis or immunotherapy agents from true disease progression. The ability to diagnose and characterize the type of glioma prior to surgical procedures could also aid neurosurgeons in tailoring the surgical approach to offer optimal benefit such as surgical planning for maximal resection when typically a diagnostic biopsy would be performed or clinical trial screening prior to initial surgical intervention which may include intraoperative surgical trials and neoadjuvant therapies. In addition, a possible advantage of detecting and characterizing the *IDH* status of aggressive gliomas rekindles the concept of neoadjuvant therapy prior to surgical resection, a frontier that has yet to be fully explored in this type of pathology. Recently, *IDH* mutation status of a solid tumor has been shown to change during tumor progression possibly due to drug response ^40^, which is only confirmed during recurrence by biopsy or resection. However, as suggested by Mazor et al ^40^, longitudinal monitoring of *IDH* status during treatment offers significant opportunities to understand the role of *IDH1* inhibitors. Last, but not least, although the cost of profiling whole-epigenome arrays may currently limit its potential application, the discovery of an alternative, sensitive and more cost-effective method to profile the epigenome of cfDNA has shown promise in several tumors^34^. Once validated, the ML approach using detected eLB may be carried out after the detection and initial treatment stages and used as real-time surveillance where frequent blood sampling could aid the treatment team in monitoring disease status, progression to a more or less aggressive phenotype, and response to specific treatment modalities. This could be done from the comfort of the physician’s office or anywhere a blood sample can be obtained rather than relying solely on diagnostic imaging or surgical excision, as these require accumulation of tissue prior to recognition of tumor activity, and offer the possibility of early intervention and preventative approaches. Serial real-time monitoring of gliomas with the relative safety of a blood test has the potential to redesign primary brain tumor diagnosis, surveillance, and therapeutic endpoints which may shed light on new opportunities that improve outcomes for patients with malignant primary brain tumors.

In summary, we developed a glioma index based on the glioma-specific CpG methylation landscape that accurately predicted the presence of these tumors. These encouraging results provide the framework to develop a blood-based epigenetic panel marker that will not only provide real-time information regarding tumor features and burden but can also be used to monitor disease progression and treatment response using a minimally invasive approach in ever-evolving tumors such as malignant gliomas.

## SUBJECTS AND METHODOLOGY

### EXPERIMENTAL DESIGN

#### Patients

We performed a retrospective study entailing archival serum and tissue from patients who received surgery to resect gliomas (N=22; composed of 15 *IDH*mut and 7 *IDH*wt; 4 *MGMT*-negative and 18 *MGMT*-positive, Extended Data Table S1) at the Henry Ford Health System (HFHS). The samples selected for this study had both serum and primary tissue available at the tumor bank of the Hermelin Brain Tumor Center (HFHS, Detroit, MI). As an early tumor tissue source site, matching tissue was submitted to TCGA from the HFHS and analyzed along with 1,122 other primary gliomas in a comprehensive pan-glioma study reported by our team ^3^. Additional serum samples from 3 non-tumor subjects were included (N=3, epilepsy). Clinical data comprised of demographic features, and pathology report (e.g., grade, histology, molecular markers, imaging, time to death, time to recurrence/progression, and treatment type) were extracted from patient medical records. The project was approved by the HFHS Institutional Review Board (IRB# 12490) and patients consented to have their specimens used for research purposes. Tissue samples were blindly reviewed by two neuropathologists (A.M. and D.C.) to determine which part of the tumor was feasible for DNA extraction (based on the percentage of necrosis, hemorrhage, infiltration, and adjacent brain, etc.) and to confirm the molecular status by immunohistochemistry and/or PCR (e.g., *IDH* and *MGMT* status). Detailed information about our cohort demographics is depicted in Table 1 and Extended Data Table S1.

#### Serum collection and processing

Peripheral blood (15 mL) was drawn for each subject into 2 BD Vacutainer SSTs (Becton Dickinson) at the time of surgery, before the opening of the dura-mater. Serum sample was separated within 1 hour from collection by centrifugation at 1,300 × g for 10 minutes at 20°C; aliquoted into up to five 2 mL cryovials and stored at −80°C until processing.

#### cfDNA isolation, quantification, and DNA methylation data generation

cfDNA was extracted from 2.2-9.3 mL aliquots of serum (Extended Data Table S1) using the Quick-cfDNA Serum & Plasma Kit according to the manufacturer protocol (Zymo Research - catalog # D4076). DNA concentration was measured with Qubit (Thermo Fisher Scientific). The concentration of cfDNA in the serum was calculated by dividing the total amount of cfDNA extracted by the amount of serum used for extraction. We then converted the concentration of cfDNA in the serum (ng/mL) into haploid genome equivalents/mL by multiplying by a factor of 303 (assuming the mass of a haploid genome 3.3 pg) ^45^.

The extracted DNA (30-300 ng) was bisulfite-converted (Zymo EZ DNA methylation Kit; Zymo Research) and profiled using an Illumina Human EPIC array (HM850K), at the USC Epigenome Center, Keck School of Medicine, University of Southern California, Los Angeles, California. The amount of bisulfite-converted DNA as well as the completeness of bisulfite conversion for each sample was assessed using a panel of MethyLight-based real-time PCR quality control assays as described previously ^58^. Bisulfite-converted DNAs were then repaired using the Illumina Restoration Kit as recommended by the manufacturer (Zymo EZ DNA methylation Kit; Zymo Research). The repaired DNA was used as a substrate for the Illumina EPIC BeadArrays, as recommended by the manufacturer and first described in Moran et al., 2016 ^59^. The raw DNA methylation data reported in this paper has been deposited to Mendeley Data (CURRENTLY embargo) at https://data.mendeley.com/datasets/cgrz6zztfg.

#### DNA methylation pre-processing

Methylation array data were processed with the minfi package in R. The raw signal intensities were extracted from the *.IDAT files and corrected for background fluorescence intensities and red-green dye-bias using the ‘noob’ function (preprocessNoob) as described by Triche et al., 2013 ^60^. The beta-values were calculated as (M/(M+U)), in which M and U refer to the (pre-processed) mean methylated and unmethylated probe signal intensities, respectively. Measurements in which the fluorescent intensity was not statistically significant above background signal (detection p value > 10^-16^) were removed from the data set. Before the analyses, we filtered out probes that were designed for sequences with known polymorphisms or probes with poor mapping quality (complete list of masked probes provided by Zhou et al. ^61^) and the X and Y chromosomes.

#### Deconvolution

To tease out the origin of the cfDNA in the serum, we applied a previously described methodology ^45^ to deconvolute the relative contribution of cell types to a given sample. Briefly, we selected 100 of the most specific hypermethylated and hypomethylated CpG probes for each cell line of interest. Given the availability of public data and the nature of our study, we included healthy cell lines and isolated cells from the blood, brain, vascular endothelial cells and adipocytes. We then used our DNA methylation derived signature and the non-negative least squares method to deconvolute our serum cohort using the standalone program provided by Moss and colleagues ^45^. We then normalized the percentages generated by the standalone program for each cell type from 0 to 100 by serum and tissue separately.

#### Supervised analysis

We performed a supervised analysis taking into account clinical-pathological attributes in both tissue or serum and used the Wilcoxon rank-sum test to identify differentially methylated sites between two groups of study (i.e., glioma vs non-glioma, *IDH*mut vs *IDH*wt) Probes were considered differentially methylated when p-values were less than 0.01 in the comparison between *IDH*mut vs *IDH*wt glioma samples (*IDH*-eLB), or 0.001 between glioma with non-tumor samples (Glioma-eLB). To identify differentially methylated probes in the serum that were tissue-specific, we calculated the differences in DNA methylation between the serum and tissue, by patient. Next we calculated the mean of the difference for each probe across glioma samples. For the *IDH*-specific analysis we calculated the mean of the difference across *IDH*mut and *IDH*wt samples. We then selected probes with less than 5% difference between tissue and serum and considered them tissue-specific.

#### Random Forest

We used a random forest ML model with the aim to classify available cfDNA methylation (serum or plasma and tissue) derived from tumor patients, patients with metastasis of unknown primary, non-tumor conditions (sepsis, pancreatic islet transplantation recipient) and non-tumor/non-diseased cell-free DNA and non-tumor brain and glioma tissue from TCGA.

To validate the performance of our ML model, for this specific approach, we randomly selected 11 glioma serum samples and used this set to compare to the 3 non-tumor serum samples and derived a new glioma-tissue specific signature (top 1000 most significant probes). We then used this set of 11 glioma serum samples and 3 non-tumor serum samples to train the random forest ML model. We tested the remaining 11 glioma serum samples using our model, ensuring to bootstrap 1000 times to reduce training/test biases. We named the output probability score of the random forest model Glioma-eLB index (GI).

#### Gene Expression and integrative analysis

First, we stratified the *IDH*-eLB CpG probes into distinct genomic regions ^41,54^ defined as “OpenSeas” (N=51 *IDH*mut, N=33 *IDH*wt), “Shores” (N=26 *IDH*mut, N=44 *IDH*wt), “Shelves” (N=10 *IDH*mut, N=4 *IDH*wt) and “CpG Islands” (N=27 *IDH*mut, N=43 *IDH*wt) (Extended Data Fig. S3E). We annotated our *IDH*-specific eLB prognostic probes (from the comparison *IDH*mut vs *IDH*wt serum samples) to their genomic location and selected probes mapped to the promoter of genes (defined as 2000 base pairs upstream and 200 base pairs downstream the TSS). We then looked into the DNA methylation levels of promoter CpGs in the corresponding glioma tissue and selected the probes in which the methylation status (i.e. hypermethylated or hypomethylated) was the same in both tissue and serum samples. Finally, we investigated the expression levels of the corresponding gene in glioma tissue and annotated their identified roles in prior cancer studies.

## ACKNOWLEDGMENTS

The authors are grateful to the patients who participated on this study. This work was supported by the Henry Ford Health System, Department of Neurosurgery and the Hermelin Brain Tumor Center Foundation. Additionally, LMP, HN, ACD, MW, and AM are supported by NCI R01CA222146 and HN, TSS, TMM, LMP, ACD, and AVC are supported by Department of Defense CA170278. We would like to thank Nancy Takacs for administrative support. We would like to thank Susan MacPhee and Kate Lawrenson for critical review of the manuscript.

## AUTHOR CONTRIBUTIONS

The contributions of the authors are as follows: Serum and tissue collection and storage: KN and AD. surgical procedure and pathology review: JS, AM, DC, AR, MR, TM, JR, IL, TW, and SK. cfDNA extraction and methylome profile: TMM, KN, and AD. Glioma-eLB and *IDH*-eLB signatures, HN, TSS, TMM, and AVC; methodology, HN, TSS, TMM, and AVC; statistical analysis, HN, TSS, TMM, and LP; pituitary tissue and serum curation, HN, TMM, MW, MM, and AVC; clinical analysis and interpretation, HN, TSS, TMM, LP, JS, KPA, TM, IL, TW, SK, and AVC; data interpretation, HN, TSS, TMM, and AVC; visualization: TSS and TMM with AVC and HN input; original draft: HN and AVC; and overall concept and coordination, HN. All authors read and approved the final manuscript.

## COMPETING INTERESTS

The authors declare to have no competing interests.

---

**Extended Data Fig. 1.**
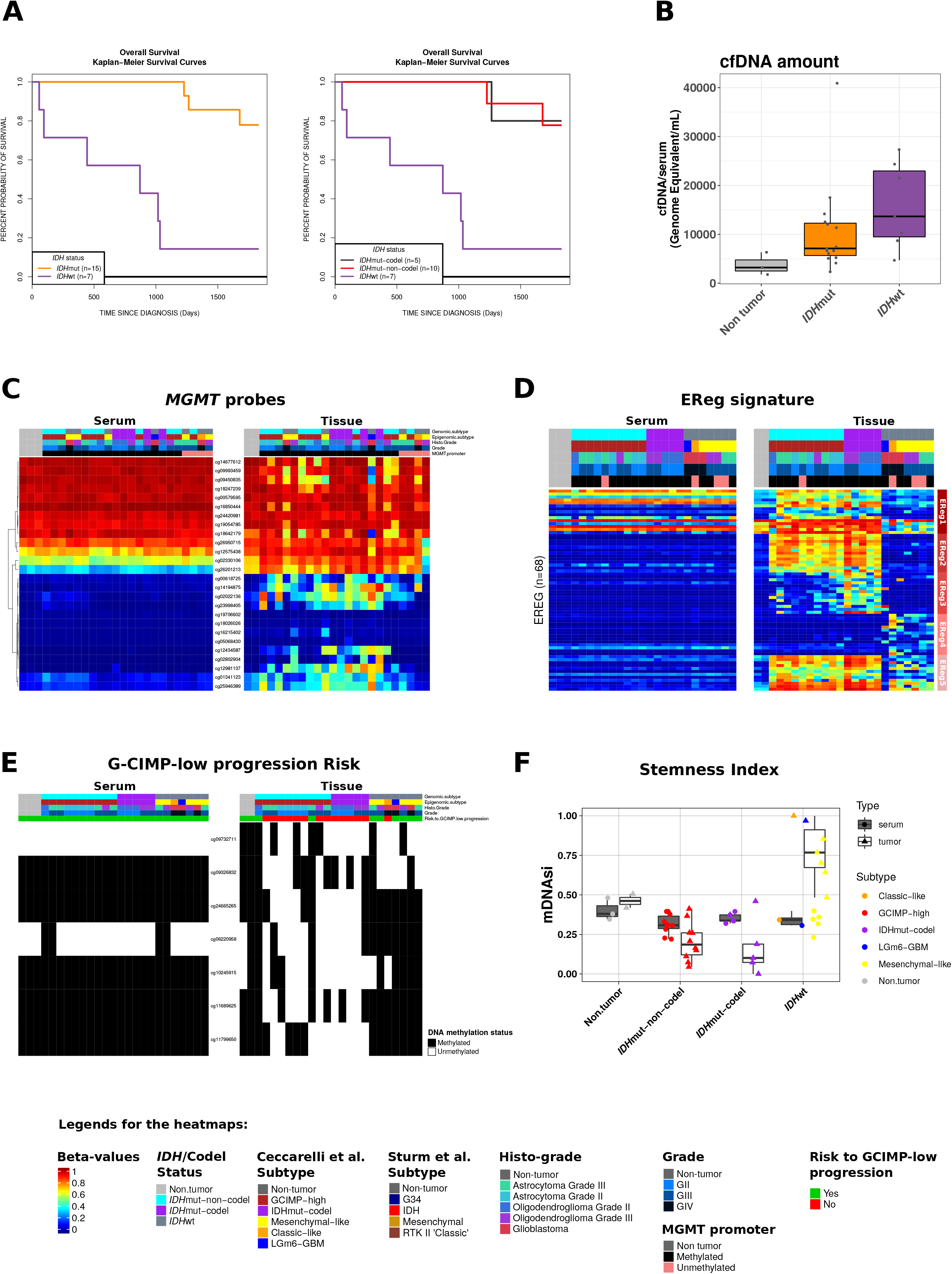
A) Kaplan-Meier survival curves showing samples separated by *IDH* status (left) and *IDH* status combined with 1p/19q co-deletion (right). Tick represents censorship. B) Total serum cfDNA concentration normalized to the genomic size (Genomic Equivalents/ml) in non-tumor, *IDH*mut and *IDH*wt samples. C) Heatmap of DNA methylation probes mapped the promoter region of *MGMT* gene. DNA methylation beta-values are represented as a color gradient from low (blue) to high (red) in C, D, and E. D) Heatmap of DNA methylation of probes that define epigenetically regulated genes in glioma subtypes. Rows represent EReg probes described by Ceccarelli et al., 2016. E) Predictive biomarkers for glioma progression. Rows represent probes that stratify gliomas into risk for aggressive recurrence (deSouza et al., 2018). Each marker was coded as white if methylated and black if unmethylated according to the published cutoffs. F) Stemness index defined by DNA methylation as described by Malta et al. 2018. Samples are stratified by genomic group and by sample type.

**Extended Data Fig. 2.**
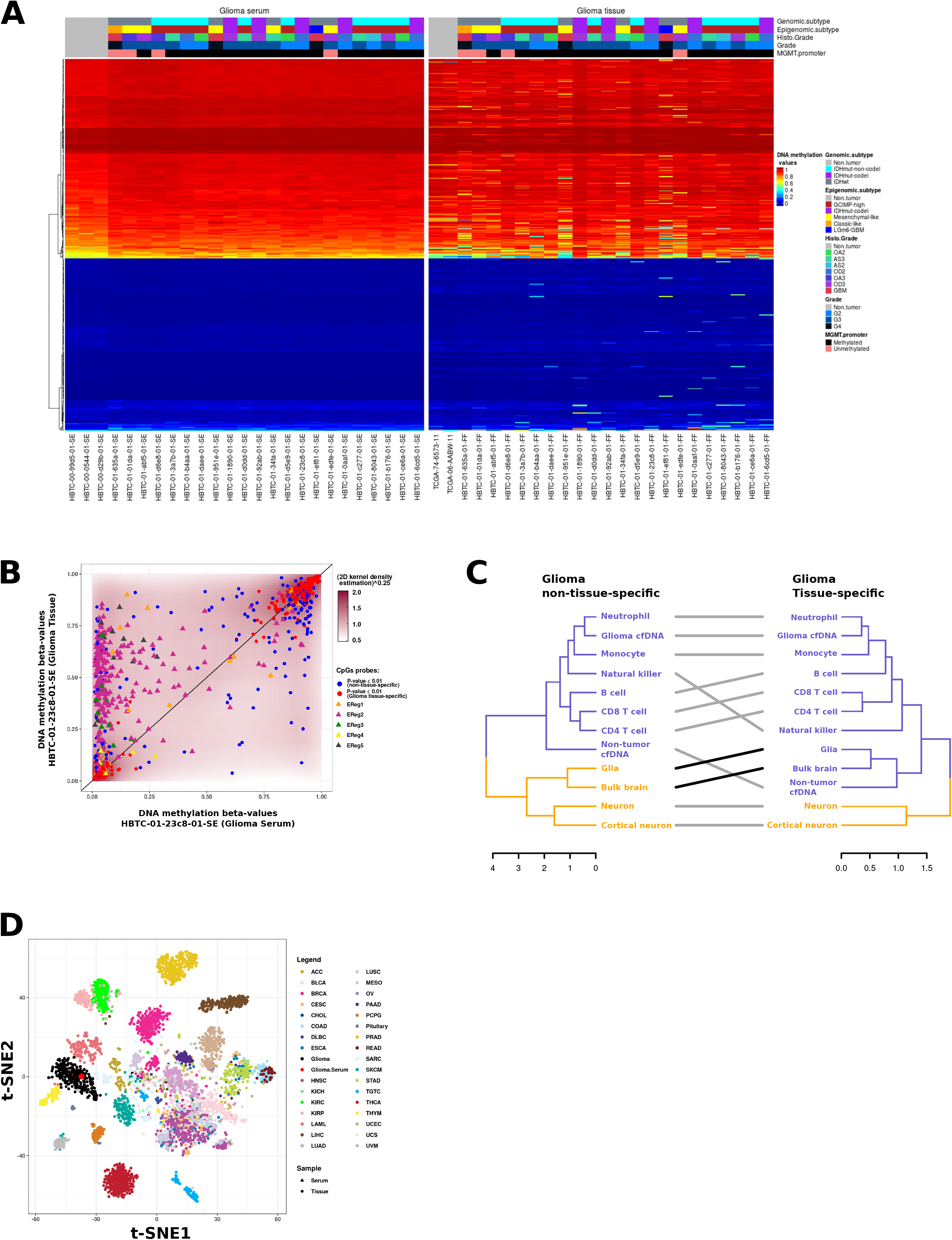
A) Heatmap of DNA methylation of Glioma-eLB probes (N = 1075 CpG sites). B) DNA methylation levels of Glioma-eLB and previously published probes (Ceccarelli et al., 2016) in the serum and in the tissue of a representative patient. C) Dendrograms of non-tumor cell types and serum using Glioma non-tissue-specific CpGs (N = 186, left) in comparison to tissue-specific (N=384, right) CpGs. D) Glioma-specific tissue-matching eLB (N=384) was used to subset the published primary tumor tissue DNA methylation (N=33 tumor types) and using t-SNE (t-distributed stochastic neighbour embedding) dimensionality reduction to visualize the similarities of each sample. As expected, each primary tumor type (circles) clusters with its known cell-of-origin. Serum cfDNA methylation of our patients cohort (triangles) clusters with the primary glioma tissue DNA methylation profiles.

**Extended Data Fig. 3.**
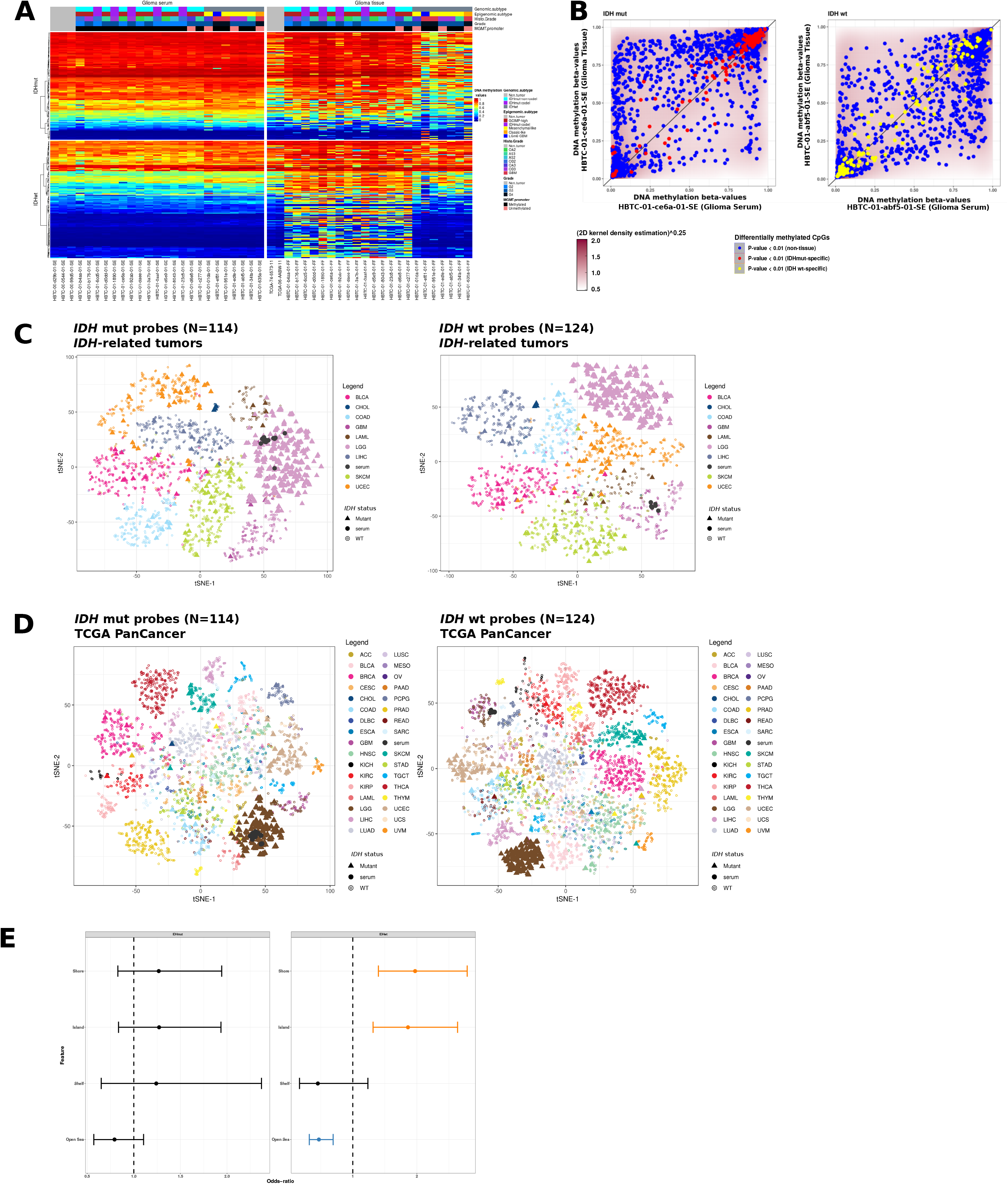
A) Heatmap of DNA methylation of *IDH-eLB* probes (N = 114 *IDH*mut-eLB and 124 *IDH*wt-eLB CpG sites). B) Comparison between DNA methylation levels of glioma tissue (y-axis) and serum (x-axis) of one representative *IDH*mut glioma patient on the left and one representative *IDH*wt glioma patient on the right. C-D) *IDH*-eLB (N=114 *IDH*mut-eLB and 124 *IDH*wt-eLB CpG sites) was used to subset the published primary tumor tissue DNA methylation data and using t-SNE (t-distributed stochastic neighbour embedding) dimensionality reduction to visualize the similarities of each sample. C) t-SNE using *IDH*mut-eLB CpGs on the left and *IDH*wt-eLB CpGs on right with primary TCGA tumor tissue from tumor types (N=9) with known *IDH* mutation. As expected, each primary tumor type (circles) clusters with its known cell-of-origin. Serum cfDNA methylation of our *IDH*mut patients cohort (triangles) clusters with the *IDH*mut primary glioma tissue and serum cfDNA methylation of our *IDH*wt patients cohort (triangles) clusters with the *IDH*wt primary glioma tissue. D) t-SNE using *IDH*mut-eLB CpGs on the left and *IDH*wt-eLB CpGs on right with primary TCGA tumor tissue (N=33 tumor types). E) Odds-ratio for the frequencies of *IDH*mut-eLB probes (left) and *IDH*wt-eLB (right), respectively, that overlap a particular molecular feature relative to the expected genome-wide distribution of the methylation platform.

**Extended Data Fig. 4:**
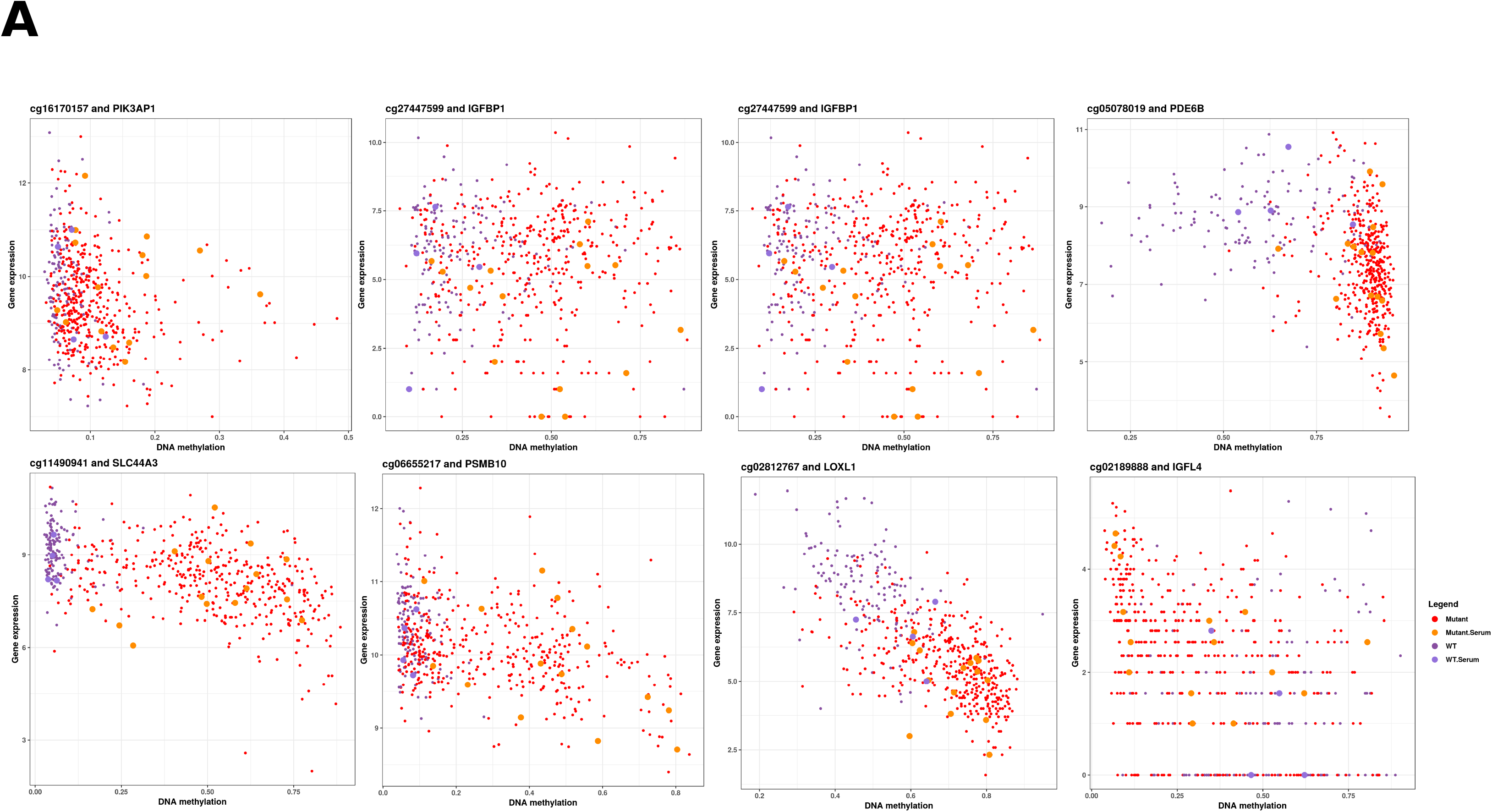
A) DNA methylation (x-axis) and expression (y-axis) scatter plot for promoter CpG associated with the corresponding gene. Each dot represents a sample. Red represents *IDH*mut glioma tissue samples, dark purple represents *IDH*wt glioma tissue samples, orange represents *IDH*mut glioma serum cfDNA and light purple *IDH*wt glioma serum cfDNA.

